# Tuning the interaction of a ParA-type ATPase with its partner separates bacterial organelle positioning from partitioning

**DOI:** 10.1101/2025.05.22.655647

**Authors:** Jordan A. Byrne, Hema M. Swasthi, Longhua Hu, Christopher A. Azaldegui, Jian Liu, Anthony G. Vecchiarelli

## Abstract

The <u>m</u>aintenance of <u>c</u>arboxysome <u>d</u>istribution (Mcd) system comprises the proteins McdA and McdB, which spatially organize carboxysomes to promote efficient carbon fixation and ensure their equal inheritance during cell division. McdA, a member of the ParA/MinD family of ATPases, forms dynamic gradients on the nucleoid that position McdB-bound carboxysomes. McdB belongs to a widespread but poorly characterized class of ParA/MinD partner proteins, and the molecular basis of its interaction with McdA remains unclear. Here, we demonstrate that the N-terminal 20 residues of *H. neapolitanus* McdB are both necessary and sufficient for interaction with McdA. Within this region, we identify three lysine residues whose individual substitution modulates McdA binding and leads to distinct carboxysome organization phenotypes. Notably, lysine 7 (K7) is critical for McdA interaction: substitutions at this site result in the formation of a single carboxysome aggregate positioned at mid-nucleoid. This phenotype contrasts with that of an McdB deletion, in which carboxysome aggregates lose their nucleoid association and become sequestered at the cell poles. These findings suggest that weakened McdA–McdB interactions are sufficient to maintain carboxysome aggregates on the nucleoid but inadequate for partitioning individual carboxysomes across it. We propose that, within the ParA/MinD family of ATPases, cargo positioning and partitioning are mechanistically separable: weak interactions with the cognate partner can mediate positioning, whereas effective partitioning requires stronger interactions capable of overcoming cargo self-association forces.

## INTRODUCTION

Bacterial microcompartments (BMCs) are protein-based organelles comprised of selectively permeable protein shells that encapsulate enzymes, thus acting as nanoscale reaction hubs that are critical to bacterial metabolism. A recent bioinformatics analysis identified 68 distinct types of BMCs across 45 bacterial phyla, indicating their widespread distribution (Sutter et al., 2021). Despite their prevalence and importance in bacterial metabolism, little is known about the spatial regulation of BMCs.

Carboxysomes are the most well-studied type of BMC. These structures encapsulate the enzyme ribulose-1,5-bisphosphate carboxylase/oxygenase (Rubisco) alongside its substrate CO[, significantly enhancing carbon fixation efficiency in autotrophic bacteria (Yeates et al., 2008). Carboxysomes contribute to an estimated 35% of global carbon fixation (Dworkin et al., 2006; Hill et al., 2020), making them a focal point for carbon capture technologies (Long et al., 2018; Flamholz et al., 2020). Additionally, carboxysomes serve as a model system for understanding BMC biology, including their spatial regulation in the cell (MacCready and Vecchiarelli, 2021).

Carboxysomes are classified into two distinct subtypes, α and β, which differ in structure, composition, and regulation (Rae et al., 2013; Kerfeld and Melnicki, 2016). While β-carboxysomes are associated with β-cyanobacteria, α-carboxysomes are found in α-cyanobacteria and certain chemoautotrophic bacteria (Sutter et al., 2021). Despite their functional similarities, α- and β-carboxysomes are phylogenetically distinct, with α-carboxysomes being more closely related to other BMC types, including those involved in bacterial pathogenesis (Kerfeld and Melnicki, 2016). Comparative studies of these subtypes have been instrumental in advancing our understanding of BMC biology and mechanisms of subcellular organization. A mechanistic understanding of carboxysome organization in the cell has come from the study of β-carboxysomes in the model cyanobacterium *Synechococcus elongatus* PCC 7942 (*S. elongatus* hereafter), and α-carboxysomes in the chemoautotroph *Halothiobacillus neapolitanus* (*H. neapolitanus* hereafter).

Spatial organization of carboxysomes within the cell is critical for inheritance upon cell division, maintaining cellular homeostasis, and optimizing carbon fixation (Savage et al., 2010; Rillema et al., 2021). This localization is tightly regulated by the two-component <u>M</u>aintenance of <u>C</u>arboxysome <u>D</u>istribution (Mcd) system (MacCready et al., 2018; MacCready et al., 2020; MacCready et al., 2021). One component, McdA, is a member of the widespread family of ParA/MinD (A/D) ATPases - found in over 96% of sequenced bacteria and position myriad cellular components (Lutkenhaus, 2012; Vecchiarelli et al., 2012, Pulianmackal et al., 2023). The partner protein, McdB, is a novel adaptor protein that bridges McdA to the carboxysome cargo (MacCready et al., 2018). In *S. elongatus*, it has been shown that McdB associates with β-carboxysomes, while McdA dimerizes in the presence of ATP and binds the nucleoid via nonspecific DNA interactions (Hakim et al., 2021). McdB stimulates McdA release from a nonspecific DNA substrate *in vitro*, which translates *in vivo* to McdA release from the nucleoid in the vicinity of McdB-bound β-carboxysomes (Hakim et al., 2021). This interplay between β-carboxysome-bound McdB and nucleoid-bound McdA results in the equidistant positioning of carboxysomes via a Brownian-ratchet mechanism (Figure 1A) (MacCready et al., 2018; MacCready et al., 2020; Hakim et al., 2021). It remains to be determined if the McdAB system functions to distribute α-carboxysomes via a similar mechanism.

**Figure 1:**
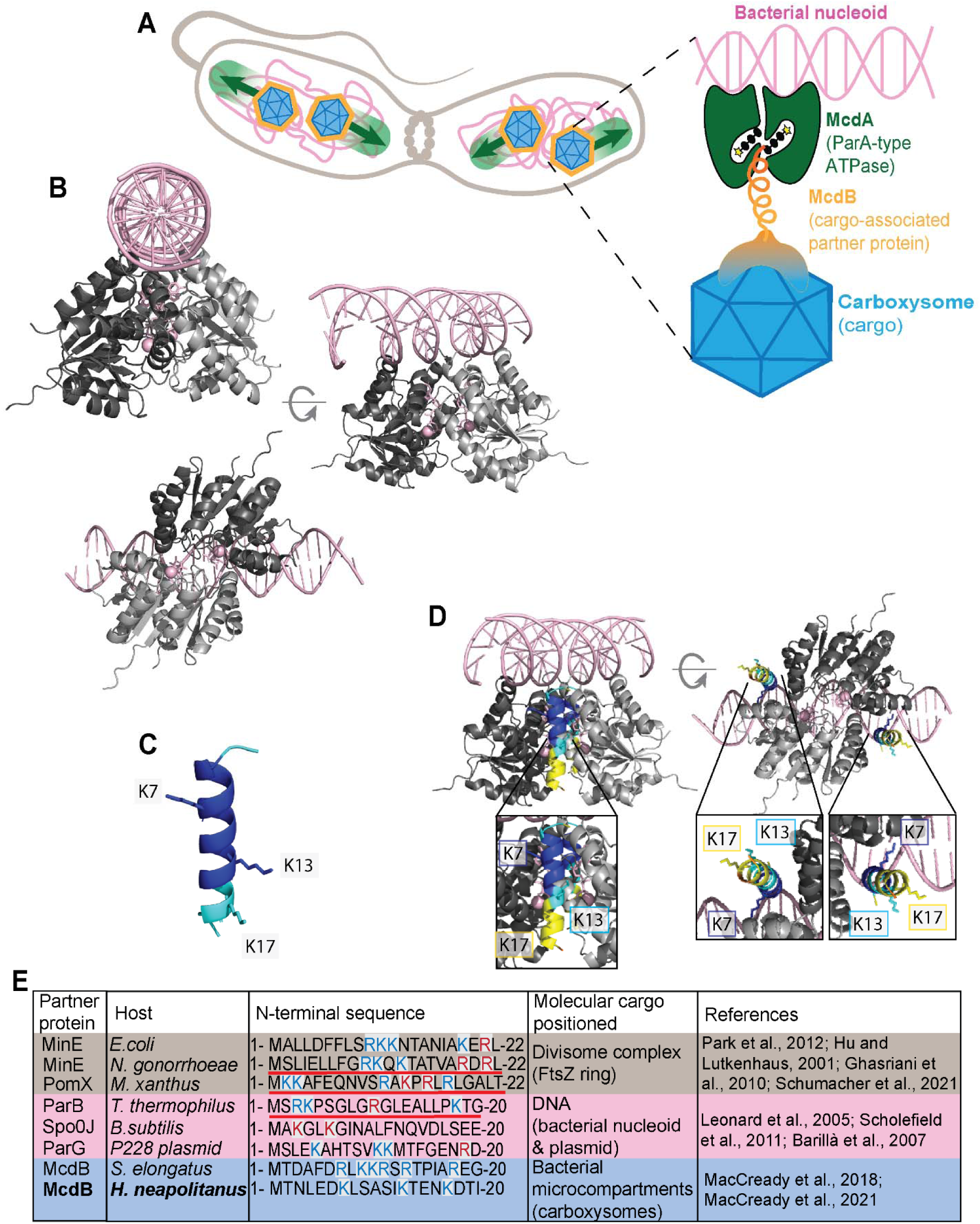
McdB is predicted to interact with McdA via basic residues at its N-terminus. **(A)** Schematic of the Brownian-ratchet model for carboxysome distribution by the McdAB system in *H. neapolitanus*. Zoom in shows the predicted quaternary DNA-McdA-McdB-carboxysome complex **(B)** AlphaFold3 (AF3) predicted structure of an ATP-bound McdA dimer (grey) docked onto a non-specific DNA substrate (pink). ATP and magnesium ions are also pink. **(C)** Predicted helical structure of the first 20 N-terminal amino acids of McdB colored by pLDDT confidence score – Dark Blue = Very high confidence (>90); Light Blue = Confident (70-90); Yellow = Low (50-70); Orange = Very low (< 50). **(D)** Two N-terminal peptides of McdB docked onto the DNA-bound McdA-ATP dimer. Peptide is colored according to pLDDT confidence score. Zoomed in boxes highlight the three lysine residues (licorice sticks) predicted for association with McdA (ipTM = 0.89). **(E)** Amino acid alignment of N-termini for the indicated ParA/MinD family partner proteins. Positively charged residues are blue and residues verified to activate their cognate ATPases are red. Underlined sequences indicate peptides that were necessary and sufficient in stimulating their cognate ATPase.

How McdB interacts with and provides specificity for its cognate McdA ATPase is unknown. While A/D ATPases display sequence, structural, and biochemical commonalities, the partner proteins linking these ATPases to their cargos are extremely diverse. Despite their diversity, data across the field supports the idea that partner proteins interact with their A/D ATPases via a positively charged N-terminus (Radnedge et al., 1998; Ravin, Rech and Lane, 2003; Barillà, Carmelo and Hayes, 2007; Ah-Seng et al., 2009; Ghasriani et al., 2010). McdB proteins share an N-terminal stretch of amino acids enriched in basic residues, but it remains to be determined if these are the specificity determinants for interaction with McdA and subsequent carboxysome distribution. Understanding these molecular details is crucial for uncovering the mechanistic basis of carboxysome positioning and for advancing our ability to engineer bacterial organelles with tailored cellular organization.

Here we show that the first 20 N-terminal amino acids of *H. neapolitanus* McdB are necessary and sufficient for interaction with McdA. We identify Lysine 7 as a critical residue in the N-terminus of McdB for McdA association. Intriguingly, substituting Lysine 7 does not fully destroy McdB interaction with McdA, and in *H. neapolitanus*, this McdB variant results in a single carboxysome cluster being positioned at mid-cell over the nucleoid. Together with mathematical simulations of our Brownian ratchet model, our data suggest weak McdA-McdB interactions can maintain the carboxysome *positioning* function of the McdAB system while losing the ability to *partition* carboxysome clusters. We propose that across the A/D family of ATPases their positioning and partitioning functions are separable - weak interactions with the cognate partner protein can be sufficient for cargo positioning, but for partitioning, the interaction and associated pulling forces must be strong enough to counteract the self-association interactions of the cargo.

## RESULTS

### The N-terminus of McdB is predicted to interact with McdA

To begin dissecting the region and residues within McdB that interact with McdA, we first performed predictive modeling and docking simulations with AlphaFold3 (AF3) (Abramson et al., 2024). Consistent with other A/D family ATPases, McdA of *H. neapolitanus* is predicted to form an ATP-sandwich dimer, with basic C-terminal residues supporting interactions with non-specific DNA (Figure 1B). Full-length McdB, on the other hand, is predicted to be monomeric and largely disordered (Figure S1A), except for the first 17 N-terminal amino acids forming an α-helix (Figure 1C). When docked onto the McdA dimer, this N-terminal α-helix is predicted to interact at the junction of the McdA dimer, both as a peptide (Figure 1D) or in the context of full length McdB (Figure S1B). An additional site for docking the N-terminal α-helix of McdB is also present on the opposing side of the McdA dimer junction (Figure 1D), suggesting one or two monomers of McdB can interact with one or both sides of an McdA dimer. An McdB variant with the α-helix removed (Figure S1C) showed no high-confidence associations with McdA (Figure S1D).

Many partner proteins are known to interact with their cognate A/D ATPase through basic residues near their N-terminus (Figure 1E) (Park et al., 2012; Hu and Lutkenhaus, 2001; Ghasriani et al., 2010; Schumacher et al., 2021; Leonard et al., 2005; Scholefield et al., 2011; Barillà, Carmelo and Hayes, 2007; MacCready et al., 2018; MacCready et al., 2021; Pulianmackal et al., 2023). The AF3 predicted N-terminal α-helix of *H. neapolitanus* McdB contains three basic residues (Lysines) at positions 7, 13, and 17 (Figure 1C). Lysines 13 and 17 of McdB are predicted to face away from the McdA dimer, whereas Lysine 7 is predicted to interact with the dimer interface of McdA (Figure 1D). Together, the predictions suggest that an N-terminal α-helix of McdB is necessary and sufficient for interaction with an ATP-bound McdA dimer, with Lysine 7 as a potentially critical residue for this association.

### The N-terminus of McdB and Lysine 7 are essential for distributing α-carboxysomes

We next set out to determine if the basic residues within the first 20 amino acids of McdB are required for distributing α-carboxysomes *in vivo*. Carboxysomes were visualized by labelling the small subunit of the encapsulated Rubisco enzyme (CbbS) with mTurquoise2 to form CbbS-mTQ (MacCready et al., 2021; Pulianmackal et al., 2023). We also performed phase-contrast imaging to monitor changes in cell morphology and the nucleoid was also imaged by DAPI-staining. Consistent with our previous work (MacCready and Tran et al., 2021; Pulianmackal et al., 2023), wildtype *H. neapolitanus* cells display carboxysomes distributed along the nucleoid (Figure 2A-C, Video 1). In cells lacking McdB (Δ*mcdB*), carboxysomes form large, nucleoid-excluded foci at one or both cell poles (Video 2). Truncation of the N-terminus, *mcdB[*Δ*2-20]*, phenocopied the McdB deletion mutant (Video 3). The data show that the N-terminus of McdB is required for distributing α-carboxysomes in *H. neapolitanus*.

**Figure 2:**
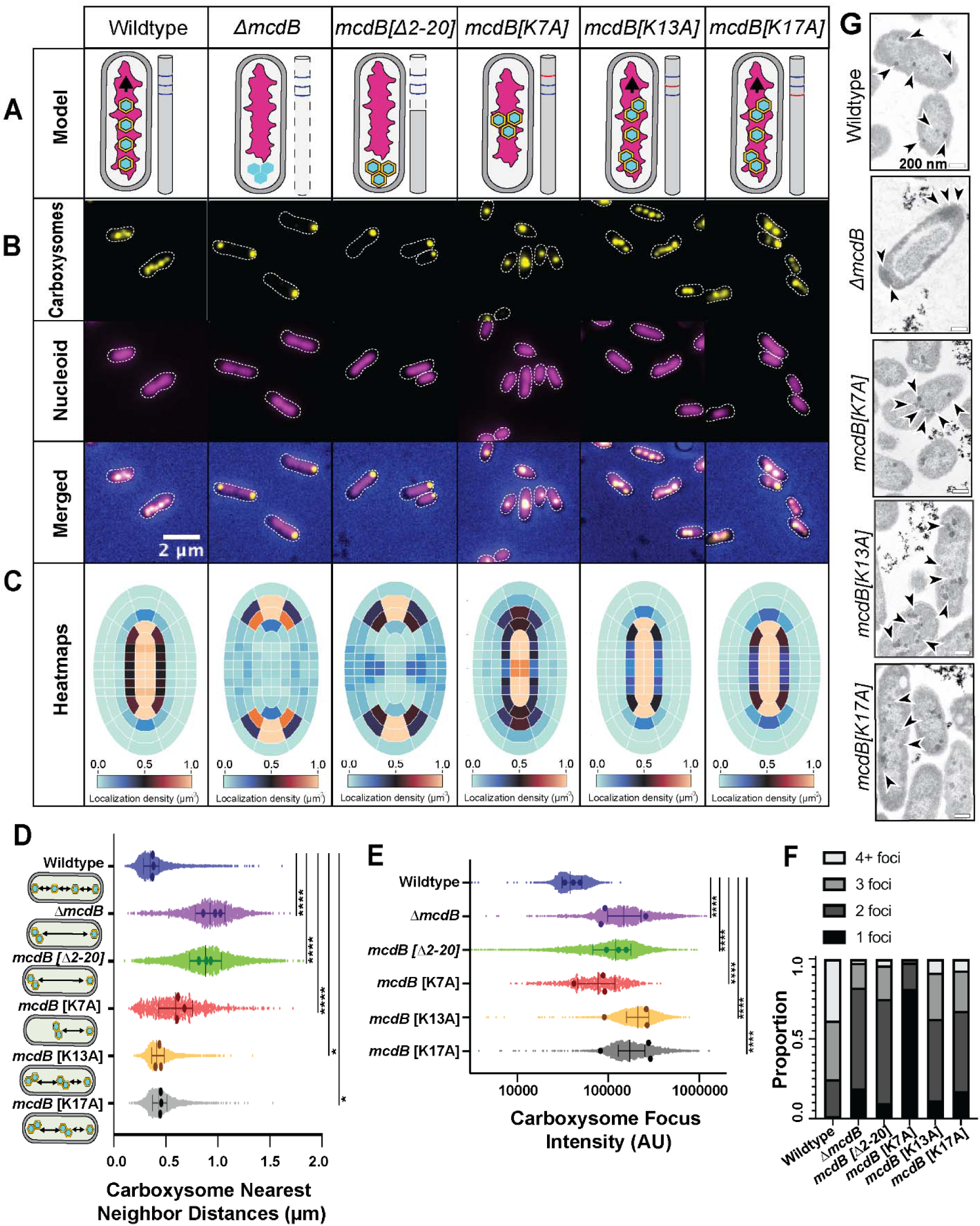
The N-terminus of McdB and Lysine 7 are essential for distributing α-carboxysomes. **(A)** Graphical representation of carboxysome localization in the indicated McdB variant strains of *H. neapolitanus*. **(B)** The carboxysome reporter CbbS-mTQ (yellow LUT) is distributed across the DAPI-stained nucleoid (magenta) in wildtype, *mcdB* [K13A], and *mcdB* [K17A] cells. Carboxysome foci localize to the cell poles in the Δ*mcdB* and *mcdB[*Δ*2-20]* strains. In the *mcdB[K7A]* strain, a single high-intensity carboxysome focus forms at mid-cell over the nucleoid. **(C)** Heatmaps display carboxysome foci distributions across the indicated cell populations. **(D)** Spacing between carboxysome foci in the same cell. Median and interquartile range are shown as bars along with averages from 3 biological replicates (dots). **(E)** As in (D), but for Carboxysome foci intensity. **(F)** Carboxysome foci number per cell. **(G)** Transmission electron micrographs of the indicated *H. neapolitanus* strains. Carboxysomes (arrows) were distributed evenly across the nucleoid in wildtype, *mcdB* [K13A], and *mcdB* [K17A] cells. Carboxysomes cluster at the cell poles in the Δ*mcdB* strain. Carboxysomes cluster over the nucleoid in the *mcdB[K7A]* strain. **(C-F)** See Table 3 for cell sample sizes for all cell population-based quantifications.

To determine if basic residues at the N-terminus of McdB play a role in distributing carboxysomes, each of the three N-terminal lysine residues was individually substituted with alanine. AF3 predicts the alanine substitutions do not interfere with the predicted helical structure of the McdB N-terminus (Figure S1E). McdB[K13A] and McdB[K17A] cells displayed carboxysome distributions similar to that of wildtype cells (Figure 2A-D, Videos 4 and 5), albeit with fewer high-intensity carboxysome foci (Figure 2E-F). The data suggest McdB[K13A] and McdB[K17A] cells have more carboxysome clustering, but these clusters are still distributed in the cell.

The majority of McdB[K7A] cells, on the other hand, displayed a singular, intense carboxysome focus (Figure 2A-F, Video 6). But unlike the McdB deletion or N-terminal truncation strains that formed high-intensity carboxysome foci at the cell poles, the carboxysome foci in McdB[K7A] cells were positioned at mid-cell. This phenotype was also observed when K7 was substituted to Glutamine or Aspartate (Figure S1F-G). Together, the data show the N-terminus of McdB, and K7 in particular, are critical for the effective distribution of α-carboxysomes in *H. neapolitanus*.

### Carboxysome clusters in McdB[K7A] cells remain associated with the nucleoid

The mid-cell positioning of a single carboxysome focus in the McdB[K7A] mutant suggested that carboxysomes were clustered but remained nucleoid associated - unlike the nucleoid-excluded clusters that form at the cell poles in the McdB deletion and N-terminal truncation strains. To determine the location of carboxysome clusters relative to the nucleoid, cells were treated with the gyrase-inhibitor ciprofloxacin to condense the nucleoid, which significantly increases the cytoplasmic space in the cell. In wildtype cells, carboxysomes remained strongly colocalized with the compacted nucleoid after ciprofloxacin treatment, despite the increased cytoplasmic space (Figure S2A), as indicated by the higher Pearson Correlation Coefficient (PCC) (Figure S2B). In McdB deletion and N-terminal truncation strains, carboxysome foci were nucleoid-excluded with a significantly reduced PCC, indicating that carboxysomes were not associated with the nucleoid (Figure S2A-B). Intriguingly, all lysine substitution mutants, including McdB[K7A], displayed carboxysomes foci that were strongly colocalized with the compacted nucleoid, with PCC values similar to that of wildtype. The data show that in the McdB[K7A] strain, carboxysome foci remain associated with the nucleoid, suggesting this McdB mutant retains some degree of McdA association, differentiating it from the McdB deletion and N-terminal truncation strains.

The small size of α[carboxysomes combined with their high copy number in the cell precludes single carboxysome resolution using traditional fluorescence microscopy. To determine if the high intensity CbbS-mTQ foci represented Rubisco aggregates, carboxysome clusters, or partitioned carboxysomes in proximity, we performed Transmission Electron Microscopy (TEM). Consistent with our interpretation of the fluorescent imaging, and as we have shown previously (MacCready et al., 2021), carboxysomes were distributed over the nucleoid region of wildtype *H. neapolitanus* cells (Figure 2G). In the absence of *mcdB*, cells displayed nucleoid-excluded carboxysome aggregates at one or both cell poles. In the McdB[K13A] and McdB[K17A] strains, carboxysomes were distributed over the nucleoid individually and as clusters. But in the McdB[K7A] strain, several assembled carboxysomes were found clustered at mid-cell, which is once again consistent with our interpretation of the fluorescence microscopy data.

### The N-terminus of McdB is necessary and sufficient for interaction with McdA

We next set out to determine if carboxysome mispositioning in the McdB mutant strains was a consequence of perturbing direct interactions with McdA using the bacterial two-hybrid (B2H) assay in *E. coli*. As shown previously (MacCready et al., 2021), McdA interacts with wildtype McdB in the B2H assay (Figure 3A). Intriguingly, a 20-amino acid N-terminal peptide of McdB was necessary and sufficient to interact with McdA at levels of association similar to that of wildtype McdB. The N-terminal truncation of McdB, on the other hand, severely abrogated the McdA association. The data show that the N-terminus of McdB is both necessary and sufficient for interaction with McdA as predicted by our AF3 modeling and docking experiments (see Figures 1 and S1).

**Figure 3:**
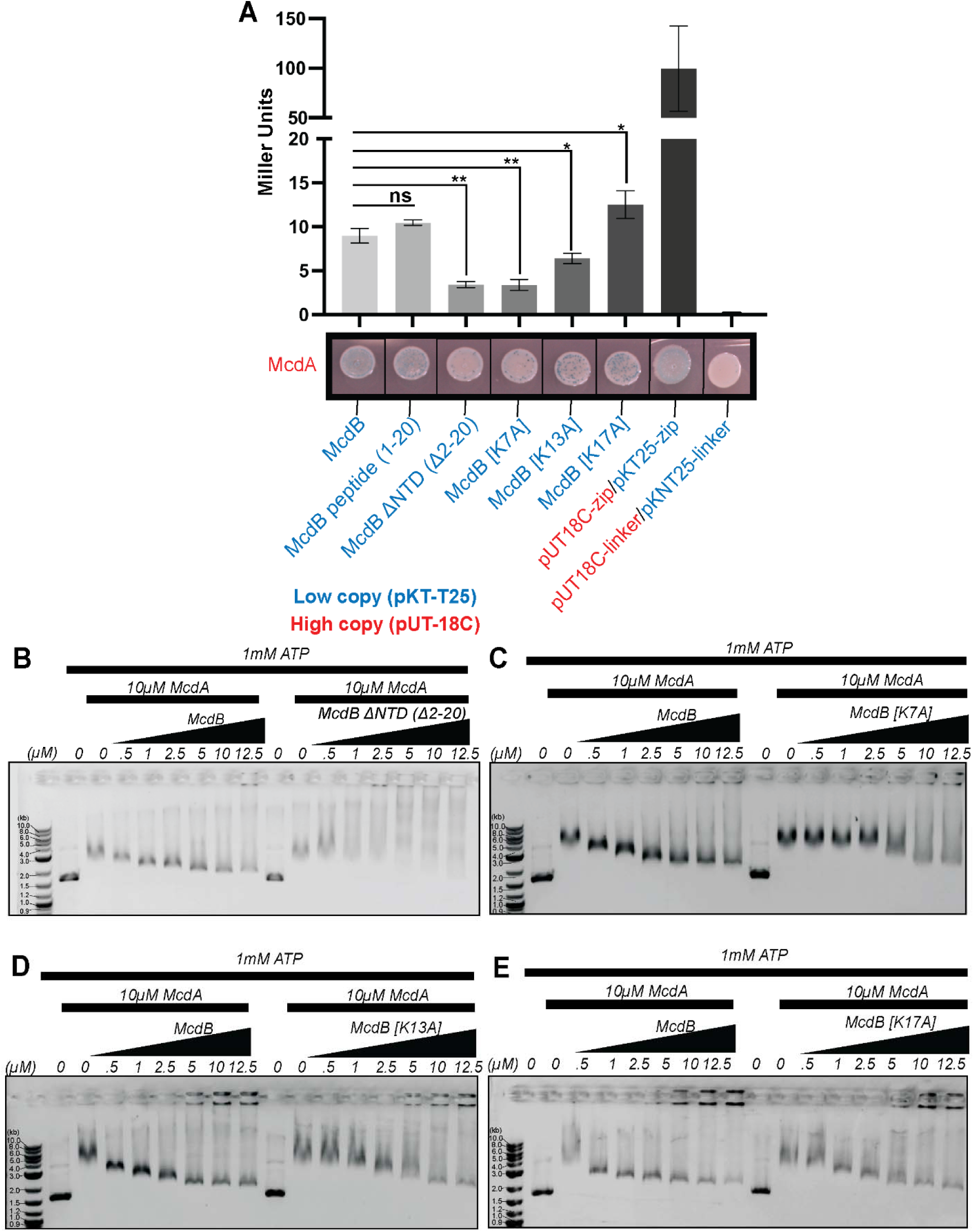
N-terminal lysine residues are required to interact with McdA and stimulate DNA release. **(A)** Bacterial 2-Hybrid (B2) interaction assay between the indicated protein pairs. The bottom image is representative of three independent experiments. Quantitative analysis of the B2H interactions (top) represent the mean and SD from three independent biological replicates. ns – not significant, *p < 0.05, **p < 0.001 by Welch’s t test. **(B)** At 10 μM McdA, increasing the WT McdB concentration stimulates McdA release from a DNA substrate (pUC19 - 2.7 kb). McdB[Δ2-20] did not stimulate release. **(C)** McdB[K7A] poorly stimulated McdA release, relative to WT McdB. **(D)** McdB[K13A] moderately stimulated McdA release, relative to WT McdB. **(E)** McdB[K17A] stimulation of McdA release from the DNA substrate was comparable to that of WT McdB.

We then assayed McdA against the N-terminal lysine substitution mutants of McdB (Figure 3A). McdB[K7A] showed little to no association with McdA, McdB[K13A] showed an intermediate level of association, and McdB[K17A] displayed the highest level of McdA association, similar to that of wildtype McdB. Together, the results reveal a tunable association strength between McdA and McdB that directly correlates with their carboxysome mispositioning phenotypes in *H. neapolitanus* cells.

### N-terminal lysine variants of McdB are defective in releasing McdA from non-specific DNA *in vitro*

We have shown previously that for the β-McdAB system of *S. elongatus*, McdB stimulates the release of McdA from a non-specific DNA (nsDNA) substrate *in vitro* (Hakim et al., 2021). *In vivo*, this activity translates to the release of McdA from the nucleoid in the vicinity of McdB-bound carboxysomes and the dynamic positioning of β-carboxysomes in the cell. Here, we performed Electrophoretic Mobility Shift Assays (EMSAs) with purified *H. neapolitanus* His-SUMO-McdA and McdB to determine if N-terminal McdB mutants were affected in their ability to stimulate McdA release from a nsDNA substrate. Consistent with our previous work on the β-McdAB system (Hakim et al., 2021), *H. neapolitanus* McdA shifted nsDNA in an ATP-dependent manner (Figure S3A). With ATP present, the nsDNA substrate exhibited decreased mobility in the presence of increasing concentrations of McdA. Conversely, when the experiment was conducted at a constant concentration of McdA (10 µM), the shift in nsDNA mobility was reversed by the addition of increasing McdB concentration (Figure S3B). McdB alone did not bind DNA (Figure S3B). Stimulation of McdA release from DNA was not observed with the N-terminal truncation mutant of McdB (Figure 3B). Importantly, purified McdB[Δ2-20] shows no degradation on an SDS-PAGE gel, showing the McdB truncation is stable (Figure S3C). Together, the findings show that for the α-McdAB system of *H. neapolitanus,* the N-terminus of McdB stimulates McdA release from DNA.

The N-terminal lysine substitution variants of McdB once again displayed a gradient of activity on McdA DNA-binding that correlated with the degree of McdA interaction in the B2H assay and with their carboxysome mispositioning phenotypes *in vivo* - McdB[K7A] poorly stimulated DNA release by McdA (Figure 3C), McdB[K13A] moderately stimulated DNA release (Figure 3D), and McdB[K17A] stimulated McdA release from DNA at levels similar to that of wildtype McdB (Figure 3E). As with wildtype McdB, the interaction between the McdB variants and McdA were ATP dependent (Figure S3D). The data confirm that the N-terminal lysine substitutions in McdB differentially impact its ability to (1) interact with ATP-bound McdA, (2) promote McdA release from non-specific DNA, and (3) distribute carboxysomes in the cell.

### Carboxysome dynamics reveal distinct organization phenotypes *in vivo*

Given the observed differences in McdA interaction and DNA-release stimulation across McdB variants, we next examined how these differences impact carboxysome dynamics *in vivo*. In wildtype cells, kymographs show that carboxysomes are highly dynamic and distributed across the nucleoid during growth and division (Figure 4A, Video 1). In contrast, cells lacking McdB exhibit restricted carboxysome diffusion, with clusters localized at the cell poles. These cells also show delayed growth and division (Figure 4A, asterisks; Video 2). In the *mcdB[K7A]* strain, a single carboxysome cluster displayed local excursions around mid-cell, but remained over the nucleoid, suggesting that McdA interactions are partially retained (Video 4). Carboxysome clusters were also dynamically positioned over the nucleoid in the *mcdB[K13A]* strain (Video 5) and to a lesser extent in the *mcdB[K17A]* strain (Video 6). But in these strains clustering was transient, with carboxysomes eventually being partitioned and distributed across the nucleoid region of the cell. These observations reveal a spectrum of positioning phenotypes that correlate with the functional tuning of the McdAB system, whereby the extent of McdA–McdB interaction directly influences carboxysome partitioning and positioning *in vivo*.

**Fig. 4.**
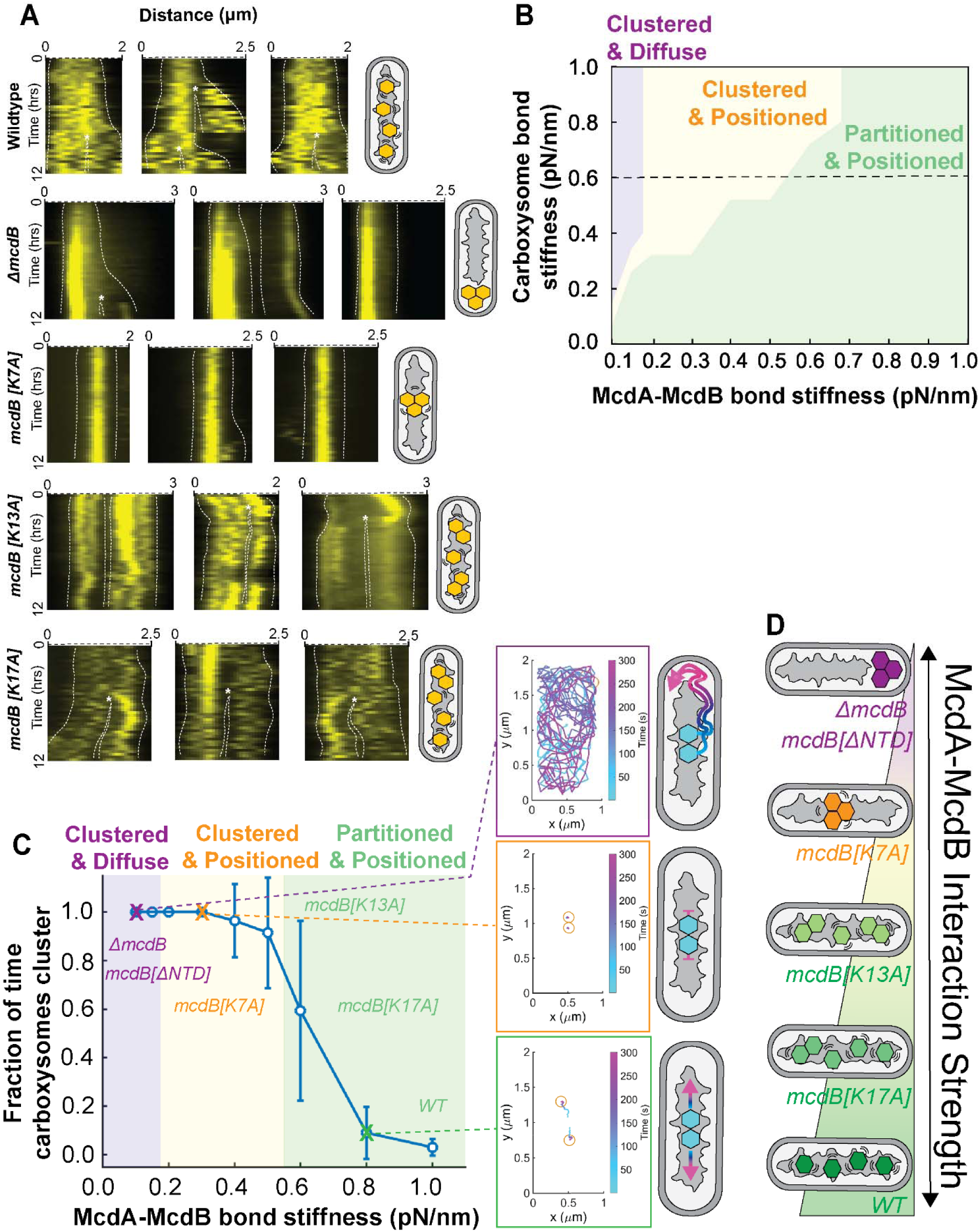
McdA-McdB binding affinities underlie distinct carboxysome behaviors *in vivo* and *in silico*. **(A)** Kymographs of carboxysome dynamics in the indicated strains over 12 hours. Dashed white lines denote cell periphery and asterisks mark division events. **(B)** Model phase diagram calculation showing how carboxysome mobility phenotypes depend on the bond stiffnesses of the carboxysome self-association interaction and the McdA-McdB interaction. For each parameter set of bond stiffnesses, we run stochastic simulations of the carboxysome system and analyze the resulting 64 trajectories. The carboxysome system is then classified into distinct phenotypes based on 1) the time fraction that the two carboxysomes stay together and 2) the motility pattern of the carboxysomes. Carboxysome mobility regimes include “clustered and diffuse” (purple), “clustered and positioned” (yellow), and “partitioned and positioned” (green). **(C)** Model predicts that the fraction of time two carboxysomes are associated with each other decreases as the McdA-McdB interaction becomes stronger. Here the spring stiffness of the carboxysome self-association was fixed (0.6 pN/nm) and the spring stiffness of McdA-McdB interactions was varied. Inset: Representative trajectories for each of the carboxysome mobility phenotypes: diffusive cluster (purple), positioned cluster (yellow), and partitioned and positioned carboxysomes (green). The trajectory color gradient denotes the evolution of the simulation time. Final carboxysome positions are marked by yellow circles. The McdA-McdB bond stiffnesses for the representative trajectories are 0.1, 0.3, and 0.8 pN/nm, respectively. **(D)** Summary model of how McdB mutants examined in this study modulate McdA–McdB interaction strength, which in turn governs the distinct *in vivo* patterns of carboxysome positioning and partitioning.

### McdA-McdB binding affinities underlie distinct carboxysome behaviors *in silico*

To assess whether differences in McdA-McdB interaction strength can explain the functional separation of carboxysome positioning and partitioning observed *in vivo*, we built upon our established Brownian ratchet model of carboxysome distribution (MacCready et al., 2018). This model is fundamentally based on the ParABS-mediated plasmid partitioning mechanism (Hu et al., 2015; Hu et al., 2017a, 2017b; Hu et al., 2021). In this study, we further developed the model by introducing self-associating interactions between carboxysome cargos, and we explored how McdA-McdB interactions counteract this self-association to coordinate carboxysome partitioning and positioning.

The stochastic simulation framework for this model is described in detail in previous work (MacCready et al., 2018; Hu et al., 2015; Hu et al., 2017a, 2017b; Hu et al., 2021). Briefly, the model treats carboxysomes as volumetrically-excluding circular disks, each 150 nm in diameter, that move along a nucleoid surface modeled as a rectangular domain. Above the nucleoid lies a cytoplasmic domain of equal dimensions. McdA molecules are modeled as volumetrically-excluding particles that bind to the nucleoid surface. After dissociation, McdA can diffuse slowly along the nucleoid or rapidly through the cytoplasm. McdB molecules are treated as fixed-position particles on the carboxysome surface. When McdB binds to nucleoid-bound McdA, the resulting McdA-McdB bond is modeled as an elastic spring subject to thermal fluctuations, which stretch the bond and generate forces that drive carboxysome motion. These elastic forces sum up to produce a net force on the carboxysome.

In line with ParABS systems and our findings here, McdB accelerates McdA dissociation from the nucleoid. When an McdA-McdB bond breaks, McdA undergoes a conformational change, leading to its rapid dissociation from the nucleoid into the cytoplasm—much faster than its intrinsic dissociation rate. Importantly, rebinding to the nucleoid is slow (Vecchiarelli et al., 2010). This mechanism causes McdA to deplete behind the carboxysome as it moves, creating an asymmetry in McdA distribution. This asymmetry biases McdA-McdB bond formation toward the carboxysome’s front, driving directional movement—hallmark behavior of a Brownian ratchet, wherein thermal motion is rectified by a self-generated McdA gradient.

To incorporate carboxysome self-association into this model, we introduced an attractive force modeled as an elastic spring between the centers of carboxysomes. The spring dynamics mimic those of the McdA-McdB interactions, but the stiffness parameters for the McdA-McdB and self-association springs are independently tunable. Our B2H data and EMSA data strongly suggest that wild-type and mutant McdB variants differ in their binding affinity for McdA (Figures 1 and 3). We therefore varied the stiffness of the McdA-McdB springs to investigate how changes in interaction strength yield different carboxysome organization phenotypes *in silico*.

We simulated a system containing two carboxysomes to explore phenotypes observed *in vivo* - such as (*i*) mispositioned clusters, (*ii*) positioned clusters, and (*iii*) partitioned and positioned carboxysomes. For each simulated trajectory, we calculated the fraction of time the two carboxysomes remained within the self-association range. Simulations ran for 300 seconds, and a pair of carboxysomes was defined as “clustered” if they were within range for > 90% of the time. To assess spatial dynamics, we measured the range of movement, defined as the maximum distance between any two points along a carboxysome’s trajectory. Clusters were considered “positioned” if their range of movement was less than 0.25 times the length of the simulation domain. Each result reflects the average of 64 simulated trajectories.

To gain a global view, we first calculated a phase diagram from our model that revealed three distinct regimes of carboxysome positioning and partitioning (Figure 4B), each of which readily explains the phenotypes observed in our *in vivo* experiments. To dissect these different motility patterns without sacrificing generality, we examined a cross-section of the phase diagram by fixing the carboxysome self-association strength (0.6 pN/nm) and varying the elastic spring constant of the McdA–McdB bonds (Figure 4C).

When the McdA–McdB interaction is too weak, it cannot separate the two carboxysomes or drive persistent motion via the Brownian ratchet mechanism. We refer to this as the “clustered and diffuse” regime (Figure 4C, purple phase and inset trajectory), where carboxysomes diffuse together across the nucleoid surface without partitioning. *In vivo*, such clusters are likely nucleoid-excluded and localize to the cell poles. At intermediate bond strength, the McdA–McdB interaction suppresses diffusion but remains too weak to split the carboxysomes. This results in the “clustered and positioned” regime (Figure 4C, orange phase and inset trajectory), in which carboxysomes remain paired and localized near mid-nucleoid. When the interaction is sufficiently strong, the carboxysomes separate and exhibit directed, persistent motion, reflecting the “partitioned and positioned” regime (Figure 4C, green phase and inset trajectory) that defines the wild-type behavior of the McdAB system.

Together, the model reveals a unifying mechanism, suggesting that the McdB mutants examined in this study modulate McdA–McdB interaction strength, which in turn governs the distinct *in vivo* patterns of carboxysome positioning and partitioning.

### N-terminal non-basic residues of McdB are also involved in McdA interaction

Up to this point, none of the single lysine substitution mutants phenocopied the Δ*mcdB* and *mcdB[*Δ*2-20]* strains, which have completely lost the ability to associate with McdA on the nucleoid and thus display nucleoid-excluded carboxysome aggregation at the cell poles. The data suggest that either the remaining lysine residues provide sufficient interactions with McdA or additional non-basic residues at the N-terminus of McdB are also playing a role in interacting with McdA. To answer this question, we made pairwise lysine-to-alanine substitutions. The *mcdB[K13A, K17A]* strain displayed distributed carboxysome foci over the nucleoid region of the cell, similar to that of the single substitution mutants (Figure S4A). However, double mutants containing the K7A mutation, *mcdB[K7A, K13A]* (Figure S4B) and *mcdB[K7A, K17A]* (Figure S4C), both displayed carboxysome clustering and nucleoid-colocalization phenotypes similar to that of *mcdB[K7A]*. This was also true for the lysine triple mutant, *mcdB[K7A, K13A, K17A]* (Figure S4D). Intriguingly, AF3 predicts the McdB[K7A, K13A, K17A] peptide retains its helical structure, but when docked onto the McdA dimer, the orientation of the peptide flips 180 degrees relative to the wildtype McdB peptide (Figure S4E). This was also true for the McdB[K7A] peptide (Figure S4F).

Together, the results further indicate that while K13 and K17 contribute to McdA interaction, K7 plays a uniquely critical role. The findings also show that non-basic residues within the N-terminus of McdB must also participate in the McdA interaction. We conclude that McdB interaction with McdA relies on a combination of key basic and non-basic residues within its N-terminal region to mediate faithful carboxysome distribution in the cell.

## DISCUSSION

Our study identifies the N-terminal 20 amino acids of *H. neapolitanus* McdB, particularly Lysine 7 (K7), as essential for mediating McdA–McdB interactions and enabling carboxysome distribution in the cell. Loss of this N-terminal region (Δ2–20), or substitution of K7, severely impairs the ability of McdB to interact with McdA, resulting in carboxysome clustering. These clusters differ phenotypically depending on the degree of McdA–McdB interaction: complete loss (ΔMcdB, Δ2–20) leads to polar aggregates, while partial loss (K7A) retains nucleoid association but fails to distribute carboxysomes evenly. These observations underscore an underappreciated but critical feature of the McdB N-terminus: it mediates not only physical binding to McdA but also determines the force balance necessary for effective spatial distribution of carboxysomes. The phenotypes observed across single-residue variants establish a gradient of positioning defects that correlate with McdA-McdB interaction strength, as confirmed by bacterial two-hybrid assays and *in vitro* EMSAs. Our work reinforces and builds upon previous studies by demonstrating the critical role of N-terminal sequences in mediating partner protein interactions with their cognate ParA/MinD ATPase (Figure 1E).

### McdA interaction via the N-terminus of McdB is likely conserved across BMC positioning systems

Our previous studies of α-McdB from *H. neapolitanus* and β-McdB from *S. elongatus* revealed striking differences both at the sequence and structural levels (MacCready et al., 2020; MacCready et al., 2021). For instance, β-McdB from *S. elongatus* is predominantly α-helical, forming a trimer-of-dimers hexamer (Basalla et al., 2023), while α-McdB from *H. neapolitanus* is largely disordered and monomeric (MacCready et al., 2021). Although α- and β-McdBs vary in secondary structure and oligomeric state, both harbor positively charged N-termini. This suggests a shared mechanism of McdA recognition, possibly through electrostatic docking at conserved interfaces. Nevertheless, subtle sequence and structural differences likely define interaction specificity and the downstream mechanics of positioning. AF3 modeling supports this idea: while the N-terminal helix of McdB reliably docks at the McdA dimer interface, mutations at K7 disrupt this interaction despite retaining helical structure predictions. Future studies will explore whether this mechanism extends to McdAB systems encoded in other BMC operons. It will also be important to experimentally determine the region and residues of McdA required for McdB association.

### Weak McdA–McdB interactions are sufficient for positioning but not partitioning carboxysomes

An unexpected yet intriguing finding is that McdB[K7A] retains enough interaction with McdA on the nucleoid to maintain mid-nucleoid positioning of carboxysome aggregates, even though it cannot separate and evenly distribute individual carboxysomes. This separation of positioning and partitioning suggests that the McdAB system operates with different interaction thresholds for these distinct functions: as illustrated in our mathematical modelling, weaker McdA–McdB interactions provide spatial confinement on the nucleoid, while stronger interactions are required to overcome carboxysome self-association and ensure their partitioning (Figure 4D). This principle likely applies broadly to other ParA/MinD-family systems, implying that positioning and partitioning are separable, tunable processes. Importantly, these results help reconcile how systems using highly divergent partner proteins can rely on conserved ATPase mechanisms while fine-tuning cargo distribution through variations in interaction strength.

### Additional N-terminal residues of McdB are involved in McdA interaction

The substitution of all lysine residues to alanine near the N-terminus of McdB still allowed for the positioning of carboxysome clusters over the nucleoid. This finding suggests that additional, yet-to-be identified, non-basic residues within the N-terminus of McdB contribute to its interaction with McdA. We previously performed *in silico* alanine-scanning mutagenesis across all N-terminal residues of *H. neapolitanus* McdB that were predicted to dock onto McdA (Pulianmackal et al., 2023). The resulting ΔΔG values predicted the extent to which each residue contributes to the stability of the McdA-McdB interaction. Notably, in addition to lysine 7 (ΔΔG = 2.4), hydrophobic residues leucine 4 (ΔΔG = 1.6), isoleucine 12 (ΔΔG = 1.8) and leucine 8 in particular (ΔΔG = 2.3) were predicted to be important for McdA interaction. Together, this *in silico* data provides a roadmap for strategic mutagenesis and the mechanistic probing of additional specificity determinants. This potential requirement for hydrophobic residues in the McdA-McdB binding interface challenges the current paradigm of partner proteins, whereby basic residues and electrostatic associations govern interaction with their cognate ATPase. Consistent with our proposal, it was recently found that for the ParAB chromosome segregation system of *Myxococcus xanthus*, conserved hydrophobic residues near the N-terminus of ParB are critical for mediating interactions with ParA (Schnabel and Osorio-Valeriano et al., 2025).

### Conclusions

Together, these results advance our understanding of ParA/MinD-based spatial organization in bacteria and lay the groundwork for engineering programmable partitioning and positioning systems for a diversity of cellular cargos. Our dissection of the molecular determinants that govern McdB interaction with McdA and the spatial organization of carboxysomes lays the groundwork for investigating how this system may be used in the design of new molecular positioning strategies. For example, now that we have shown the first 20 amino acids of McdB are both necessary and sufficient for interaction with McdA, we are leveraging this minimal motif in developing molecular tagging systems with potential applications in synthetic biology and biotechnology. Our findings also provide broader insights into the diversity of ParA/MinD-mediated cargo organization across bacterial species. Specifically, we demonstrated how subtle tuning of the interaction between a cargo adaptor and its ParA/MinD ATPase partner can distinctly regulate spatial positioning. In *H. neapolitanus*, the N-terminus of McdB - particularly residue K7 - acts as a critical determinant of carboxysome positioning. Disrupting this interaction weakens cargo partitioning while preserving mid-nucleoid localization, suggesting that these two functions are mechanistically separable across the ParA/MinD family of positioning systems.

## MATERIALS AND METHODS

### Strain List

All strains described in this study (see Supplemental Table 1) were constructed using the CbbS-mTQ carboxysome reporter strain of *Halothiobacillus neapolitanus* (Parker) Kelly and Wood (ATCC® 23641™).

### Protein Structure Prediction

All structural predictions and docking models were generated using the AlphaFold3 (AF3) server (Abramson et al., 2024). For each McdA docking prediction with McdB variants, we generated five structures with the default hyperparameters. We selected docked models based on the ipTM scores of the binding interface residues and their similarity to previously resolved ParA-like ATPase/partner-protein crystal structures. Structures were visualized and colored with PyMOL (Version 3.1.4.1).

Interface-predicted template modelling (ipTM) indicates the accuracy of the estimated interface between subunits in the multimer. An ipTM score higher than 0.8 represents high-quality predictions, while values below 0.6 suggest an inaccurate or failed prediction. Predications with ipTM values between 0.6 and 0.8 are ambiguous, meaning the structure could be correct or incorrect. All positive McdA-McdB docking predictions presented here had ipTM scores higher than 0.8.

The predicted local distance difference test (pLDDT) is a metric used by AF3 to estimate local confidence at the level of individual amino acid residues in a predicted protein structure. It is based on the local distance difference test; it does not rely on superposition but assesses the correctness of the local distances and provides a score of how well the prediction would agree with an experimental structure. pLDDT scoring is represented through gradient coloring, scaled from 0 to 100, with higher scores indicating higher confidence. A pLDDT above 90 (dark blue) is the highest confidence category, in which both backbone and side chains are predicted with the most accuracy. pLDDT above 70 (light blue) usually corresponds to a correct backbone prediction with the misplacement of some side chains. pLDDT confidence scores for all AF3 models are specified in figure legends.

### Construct Design and Cloning

Point mutations were synthesized via PCR and verified by sequencing. Constructs contain flanking DNA ranging from 750 to 1100 bp in length upstream and downstream of the targeted insertion site to promote homologous recombination into target genomic loci. Plasmid isolation was performed using chemically competent *E. coli* Top10 cells. A minimum of 200 ng of isolated plasmid was incubated with competent *H. neapolitanus* cells for 5 minutes on ice. This mixture was then transferred to a tube containing 4 mL ice-cold S6 medium without antibiotics and incubated on ice for 5 min. Transformations were recovered for 18 hours, while shaken at 130 RPM, at 30 °C, in air supplemented with 5% CO_2_. Clones were selected by plating on a selective medium with antibiotics. Colonies were restreaked on selective plates until resultant colonies were successfully verified for gene insertion and absence of vector via colony PCR. Sequence of the inserted gene mutation was verified through Illumina sequencing.

### Media and Growth Conditions

*H. neapolitanus* cultures were grown in ATCC® Medium 290: S6 medium for Thiobacilli (Hutchinson et al., 1965) and incubated at 30 °C, while shaken at 130 RPM in air supplemented with 5% CO_2_. Strains were preserved frozen at −80 °C in 10% DMSO.

### H. neapolitanus Competent Cells

Competent cells of *H. neapolitanus* were made by inoculating 10 mL S6 medium with a single colony. After 40 hours of growth, 10 mL inoculum was added to 1[L of S6 medium and grown for 18 hours to an OD600 of 0.1. Cells were separated into two 500 mL aliquots and harvested via centrifugation at 5000[x[g for 20 min at 4 °C. Pellets from each 500 mL aliquot were washed first with 250 mL, then 25 mL of ice-cold nanopore water. All wash centrifugation steps were performed at 3000xg for 30[min at 4 °C. The resulting pellet after washing was resuspended in 200 µL of ice-cold nanopore water. Competent cells were used immediately or frozen at −80 °C for future use. Frozen competent cells were thawed at 4 °C before use.

### Fluorescence Microscopy

All live-cell microscopy was performed using exponentially growing cells (OD600 .05-.1). 2 µL of cells were dropped onto a pad of S6 medium + 2% UltraPure agarose (Invitrogen, catalog number 16500)[and imaged on a 35-mm glass-bottom dish (MatTek, catalog number P35G-1.5-14-C). All fluorescence and phase contrast imaging were performed using a Nikon Ti2-E motorized inverted microscope controlled by NIS Elements software with a SOLA 365 LED light source, a 100X Objective lens (Oil CFI Plan Apochromat DM Lambda Series for Phase Contrast), and a Photometrics Prime 95B Back-illuminated sCMOS camera or a Hamamatsu Orca Flash 4.0 LT + sCMOS camera.

For the native CbbS fluorescent fusion, the sequences encoding the fluorescent protein *mTurquoise* (mTq) were attached to the 3′ region of the native *cbbS* coding sequence, separated by a GSGSGS linker. A kanamycin resistance cassette was inserted immediately downstream of the *cbbS* coding sequence. The mutant was selected by plating on S6 agar plates supplemented with 50 µg/ml of kanamycin. The CbbS-mTQ fusion was verified by PCR.

CbbS-mTQ labeled carboxysomes were imaged using a “CFP” filter set (C-FL CFP, Hard Coat, High Signal-to-Noise, Zero Shift, Excitation: 436/20 nm [426-446 nm], Emission: 480/40 nm [460-500[nm], Dichroic Mirror: 455 nm). DAPI fluorescence was imaged using a standard “DAPI” filter set (C-FL DAPI, Hard Coat, High Signal-to-Noise, Zero Shift, Excitation: 350/50 nm [325-375 nm], Emission: 460/50 nm [435-485 nm], Dichroic Mirror: 400 nm).

### Timelapse Microscopy

For multigenerational long-term microscopy, 2 µL of exponentially growing cells were dropped onto a pad of S6 medium + 1.5% UltraPure agarose (Invitrogen, catalog number 16500)[and imaged on a 35-mm glass-bottom dish (MatTek, catalog number P35G-1.5-14-C). Cells were preincubated in dishes at 30 °C in 5% CO_2_ for at least 30 minutes before image acquisition. Temperature, humidity, and CO_2_ concentrations were controlled with a Tokai Hit Incubation System. NIS Elements software was used for image acquisition. Videos were taken at one frame per 15 min for a duration of 18 hours.

### Nucleoid (DAPI) Staining for Live Imaging

Cells were harvested by centrifugation at 5000xg for 5 min. Following centrifugation, the cells were washed in PBS, pH 7.4. The resulting cell pellet was resuspended in 100 μL PBS and stained with DAPI (<u>Invitrogen, catalog number S34862</u>) at a final concentration of 25 µM. The samples were incubated in the dark, at room temperature for 10-15 min immediately before imaging. Stained cells were pelleted and resuspended with 100 μL PBS before loading onto an S6 medium agarose pad.

### Ciprofloxacin Treatment

Cells were collected at an OD of 0.05–0.1 and treated with 50 µM ciprofloxacin for 24 hr. Cells were then stained with a final concentration of 25 µM DAPI for 15 min in the dark and imaged.

### Image Analysis

#### Cell segmentation

Phase contrast images were used for cell segmentation via the Omnipose (Cutler et al., 2022) package in Python. A custom Python script was implemented for batch segmentation: PhaseMasks_omni.py. Prior to segmentation, a Gaussian blur (SD of Gaussian = 1 pixel; skimage.filters.gaussian) was applied to the images. The bact_phase pre-trained model was used for segmentation. Cells touching the borders of the image were ignored, segmented objects smaller than 150 pixels were removed, and erroneous segmentations were manually corrected using the Omnipose GUI or excluded from further analysis if cell boundaries were not clear from the phase image. Resulting phase masks were used for subsequent single-cell analyses.

#### Carboxysome focus detection

Carboxysome foci were detected using a custom Python script: detect_carboxysomes.py: first, the phase masks were used to individually analyze every cell in the field of view. The fluorescent signal in each cell was then sharpened using the scikit-image (van der Walt et al., 2014) function skimage.filters.unsharp_mask; this step was followed by a Gaussian blur (skimage.filters.gaussian). Using this preprocessed image, carboxysome foci were detected with the Laplacian of Gaussian (LoG) algorithm (skimage.feature.blob_log). See Supplemental Table 2 for parameters used. Putative foci signals were fit to a 2D Gaussian for subpixel localization. Nearest neighbor distances between adjacent carboxysome foci was determined using the custom function nearest_neighbor. Fluorescent focus intensities were determined by creating a circular mask centered on the peak of the Gaussian fit with a radius of 1.5x the standard deviation of Gaussian fit. The total intensity of the focus is the sum of pixel intensities in the circular mask. See Supplemental Table 3 for cell sample sizes used in the analysis.

#### Carboxysome localization heatmaps

The localization coordinates of detected carboxysome foci and cell masks were used to make localization density heatmaps with the spideymaps tool (https://github.com/BiteenMatlab/spideymaps) in Python described previously in Dudley et al., 2025 [ADD REF for Claire’s paper].

### Bacterial Two-Hybrid and β-galactosidase Activity Assay

N-terminal T18 and T25 fusions of McdA and all McdB mutant variants were constructed using the plasmids pKT25 and pUT18C. Plasmids were sequence-verified and co-transformed into *E. coli* BTH101 in both pairwise combinations (Karimova *et al*., 1998). 35 colonies of T18/T25 cotransformants were cultured in LB medium with 100 μg/ml ampicillin, 50 μg/ml kanamycin and 0.5 mM IPTG overnight at 30 □C with 225 rpm shaking. Overnight cultures were spotted on indicator LB X-gal plates supplemented with 100 μg/ml ampicillin, 50 μg/ml kanamycin and 0.5 mM IPTG. Plates were incubated in the dark at 30°C at least 24 hours before imaging. To quantify the interactions between hybrid proteins (Miller 1972), β-galactosidase activity measurements were performed as previously described (Battesti and Bouveret 2012) with slight modifications. Two hundred microliters of the overnight cultures were transferred into glass tubes containing 800 μl of Z buffer (45 mM Na_2_HPO_4_.12H_2_O, 45 mM NaH_2_PO_4_.H2O, 10 mM KCl, 1 mM MgSO_4_.7H_2_O, 38.5 mM β-mercaptoethanol). A drop of 0.01% SDS and two drops of chloroform were added, followed by 15 s of vortexing to facilitate cell permeabilization. Once chloroform settled to the bottom (∼ 15 s after mixing), 50 µl of the reaction mix were transferred into a 96-well flat-bottom microplate filled with 150 µl Z buffer already pre-equilibrated at 28°C in a SpectraMax® iD3 microplate reader (Molecular Devices). To start the reaction, 40 µl 0.4% *o*-nitrophenyl-β-d-galactoside (ONPG) was added and measurements at OD_420_ were taken every 2 mins for 1 hour at 28°C using the microplate reader. Concurrently, 50 µl of the overnight cultures were added to a well plate containing 150 µl LB for OD_600_ measurement in the microplate reader. β-galactosidase enzymatic activities, in Miller Units (MU), were calculated using the formula: MU = A_420_/(incubation time in minutes × culture volume in milliliters × OD_600_).

### Expression and Purification of His-SUMO-tagged McdA

McdA with an N-terminal His-SUMO tag was overexpressed in BL21(DE3) cells using the pET15b vector. An overnight culture grown in LB medium supplemented with 100 μg/mL carbenicillin was diluted 1:100 into fresh LB medium. The culture was grown at 37 °C until an OD600 of 0.4–0.6 and protein expression was induced with 100 μM IPTG. Induction proceeded for 16 hours at 16 °C. The harvested cell pellets were stored at -80 °C. For purification, the cell pellet from 500 mL of induced culture was resuspended in 30 mL of lysis buffer (50 mM Tris-HCl, 1 M KCl, pH 8.0, 10% glycerol, 5 mM β-mercaptoethanol (BME), and one protease inhibitor tablet (Thermo Fisher)). Lysozyme (30 mg) was added, and the suspension was incubated on ice for 10 minutes. Cells were lysed by sonication using a 10-second pulse-on/20-second pulse-off cycle at 50% power for 10 minutes. The lysate was cleared by centrifugation at 15,000g for 30 minutes at 4 °C, followed by filtration through a 0.45 μm filter. The clarified lysate was applied to a 5 mL HisTrap HP column (Cytiva) pre-equilibrated with Buffer A (50 mM Tris-HCl, 1 M KCl, pH 8.0, 10% glycerol, 5 mM BME). The column was washed with five column volumes of Buffer A containing 5% Buffer B (50 mM Tris-HCl, 1 M KCl, 500 mM imidazole, pH 8.0, 10% glycerol, 5 mM BME). The protein was eluted using a 5–100% gradient of Buffer B on an ÄKTA Pure system. Eluted fractions containing McdA were pooled and further purified by size-exclusion chromatography using a HiLoad 16/600 Superdex 200 pg column (GE Healthcare) equilibrated in Buffer A. Peak fractions were concentrated, filtered through a 0.45 μm filter, and stored at -80 °C.

### Expression and Purification of McdBs

McdB or mutants with an N-terminal SUMO tag was overexpressed in E. coli BL21 (AE) cells using the pET11b vector. LB medium supplemented with 100 μg/mL carbenicillin was inoculated with 1% (v/v) overnight culture and incubated at 37 °C with shaking. Protein expression was induced at an OD600 of 0.4–0.6 by adding 1 mM IPTG, followed by incubation for 4 hours at 37 °C. Induced cells were harvested by centrifugation and stored at -80 °C until further use. For purification, the cell pellet (from 1 L) was resuspended in 30 mL of Buffer A (50 mM Tris-HCl, 300 mM KCl, pH 8.4, 5% glycerol, 5 mM BME) supplemented with one tablet of protease inhibitor. Lysozyme (30 mg) was added to the suspension, followed by immediate sonication at 50% power using a 10-second pulse-on and 20-second pulse-off cycle for a total of 7 minutes. The lysate was clarified by centrifugation at 15,000 × g for 30 minutes and then filtered through a 0.45 μm membrane filter. The clarified lysate was loaded onto a 5 mL HiTrap column and washed with five column volumes of Buffer A with 5% Buffer B (50 mM Tris-HCl, 300 mM KCl, 500 mM imidazole, pH 8.4, 5% glycerol, 5 mM BME). Protein was eluted using a 5–100% gradient of Buffer B on an ÄKTA Pure system. Eluted fractions containing His-SUMO-McdB were pooled and concentrated. The concentrated protein solution was diluted to achieve a final imidazole concentration < 100 mM. Ulp1 protease was added at a 1:100 ratio (enzyme: protein) and incubated at room temperature with gentle shaking to cleave the His-SUMO tag from McdB. The cleaved McdB was further purified by size-exclusion chromatography using a HiLoad 16/600 Superdex 75 pg column (GE Healthcare) equilibrated in Buffer A. Peak fractions were concentrated and stored at -80 °C.

### Electrophoretic Mobility Shift Assay (EMSA) DNA Binding Assay

EMSA was performed in a final reaction volume of 10 μL containing 50 mM Tris (pH 8.0), 1 M KCl, 10% glycerol, 5 mM BME, and 5 mM MgCl2. The DNA substrate used was the pUC19 plasmid (10 nM). The reaction mixture included SUMO-McdA, 1 mM ATP or 1 mM ADP, DNA (10 nM), and McdB or mutant proteins in the presence of 1 mM ATP or 1 mM ADP. Reactions were incubated for 30 minutes at 23°C. After incubation, 1 μL of DNA loading dye was added to each sample. The samples were then loaded onto a 1% agarose gel prepared in TAE buffer with ethidium bromide. Electrophoresis was carried out at 100 V for 60 minutes. Following electrophoresis, the gel was visualized using a Li-Cor Odyssey imager.

### Transmission Electron Microscopy

30 mL of *H. neapolitanus* log cultures were pelleted and fixed overnight with 3% Glutaraldehyde and 3% Paraformaldehyde in 0.1 M Cacodylate Buffer, pH 7.2 at 4°C. After 3 washes with 0.1 M Cacodylate Buffer, cells were post-fixed for 1 hour in 1.5% K4Fe(CN)6 and 2% OsO4 in 0.1 M Cacodylate Buffer at room temperature. After cells were washed 3 times with 0.1 M Cacodylate Buffer, cells were then washed 3 more times with 0.1 M Na2 in Acetate Buffer, pH 5.2 at room temperature. Cells were immobilized in 1% agarose and sliced into 70 nm sections. Sections were en bloc stained for 1 hour with 2% Uranyl Acetate in 0.1 M Na2 and Acetate Buffer, pH 5.2 at room temperature. After sections were washed 2 times with 0.1 M Na2 in Acetate Buffer, pH 5.2, sections were washed 1 time with Milli-Q Water. Slices were dehydrated in an increasing series of ethanol (30%, 50%, 70%, 80%, 90%, 95%) washes for 15 min each at 4°C. Slices were dehydrated with 100% ethanol twice, then acetone once for 15 minutes each at room temperature. Slices were infiltrated with a ratio of acetone:Spurr’s Resin, 2:1 for 1 hour, 1:1 for 2 hours, then 1:2 for 16 hours at room temperature. A final incubation in full strength resin was performed for 24 hours at room temperature under vacuum. Cells were then embedded in blocks, labeled, and incubated under vacuum for 30 minutes at room temperature, before being transferred to a 70°C oven to polymerize for 24 hours. Thin sections of approximately 70 nm were obtained with a Leica UC7 Ultramicrotome. Sections were visualized on a JEOL 1400-plus transmission electron microscope equipped with an XR401 AMT sCMOS camera.

### Statistical Analysis

For whole population analyses, such as fluorescence intensity and cell length measurements, we performed a non-parametric Wilcoxon (Kruskal-Wallis) test followed by Dunn’s multiple comparisons test. For percent of population analyses, such as foci number, populations were analyzed as separate field of views and the summary values were plotted as individual data points. A Brown-Forsythe and Welch ANOVA test was performed, followed by Dunnett’s T3 multiple comparisons test. GraphPad Prism (version 10.4.2) was used to perform all analyses.

### Native Fluorescent Fusions

For strains that contain native CbbS fluorescent fusions, the sequences encoding the fluorescent protein mTurquoise (mTQ) were attached to the 3[ region of the native cbbS coding sequence, separated by a GSGSGS linker. A kanamycin resistance cassette was inserted immediately downstream of the cbbS coding sequence. The CbbS-mTQ fusion was verified for insertion and absence of vector via colony PCR. Sequence was verified through Illumina sequencing.

## Abbreviations

BMC: Bacterial Microcompartment
mNG: monomeric NeonGreen
mTQ: monomeric Turquoise2
McdA: Maintenance of Carboxysome Distribution protein A
McdB: Maintenance of Carboxysome Distribution protein B
ParA: Partition protein A
ParB: Partition protein B
ATP: Adenosine triphosphate
nsDNA: non-specific DNA
DAPI: 4′,6-diamidino-2-phenylindole

## ACKNOWLEDGEMENTS

We would like to thank members of the Vecchiarelli Lab for providing feedback and helpful discussions, and to Vayda Nyland for assistance with the Bacterial-Two-Hybrid assay. We also thank Megan Bennett from the University of Michigan Microscopy Core for assistance with TEM grid preparation. This work was supported by funding from the National Science Foundation to A.G.V. (Grant no. 1941966), the National Institute of General Medical Sciences of the National Institutes of Health to A.G.V. (Grant no.532 R35GM152128), the NIH Cellular Biotechnology Training Program grant to J.A.B. (T32 GM145304), and a Rackham Graduate Research Grant (127751) to J.A.B. J. L. was supported by start-up funds from the Johns Hopkins University School of Medicine, Johns Hopkins Catalyst award, the National Science Foundation (2105837 and 2148534), and NIH R01 GM148459.

## AUTHOR CONTRIBUTIONS

Conceptualization, J.A.B. and A.G.V.; Investigation, J.A.B. and H.M.S.; Software and Image Analysis, J.A.B. and C.A.A.; Mathematical Modelling, L.H. and J.L. Supervision, A.G.V. and J.L.; Writing – original draft, J.A.B. and A.G.V.; Writing – review & editing, J.A.B., H.M.S., C.A.A., L.H., J. L. and A.G.V.

