## Supplementary Information for "Tuning the interaction of a ParA-type ATPase with its partner separates bacterial organelle positioning from partitioning"

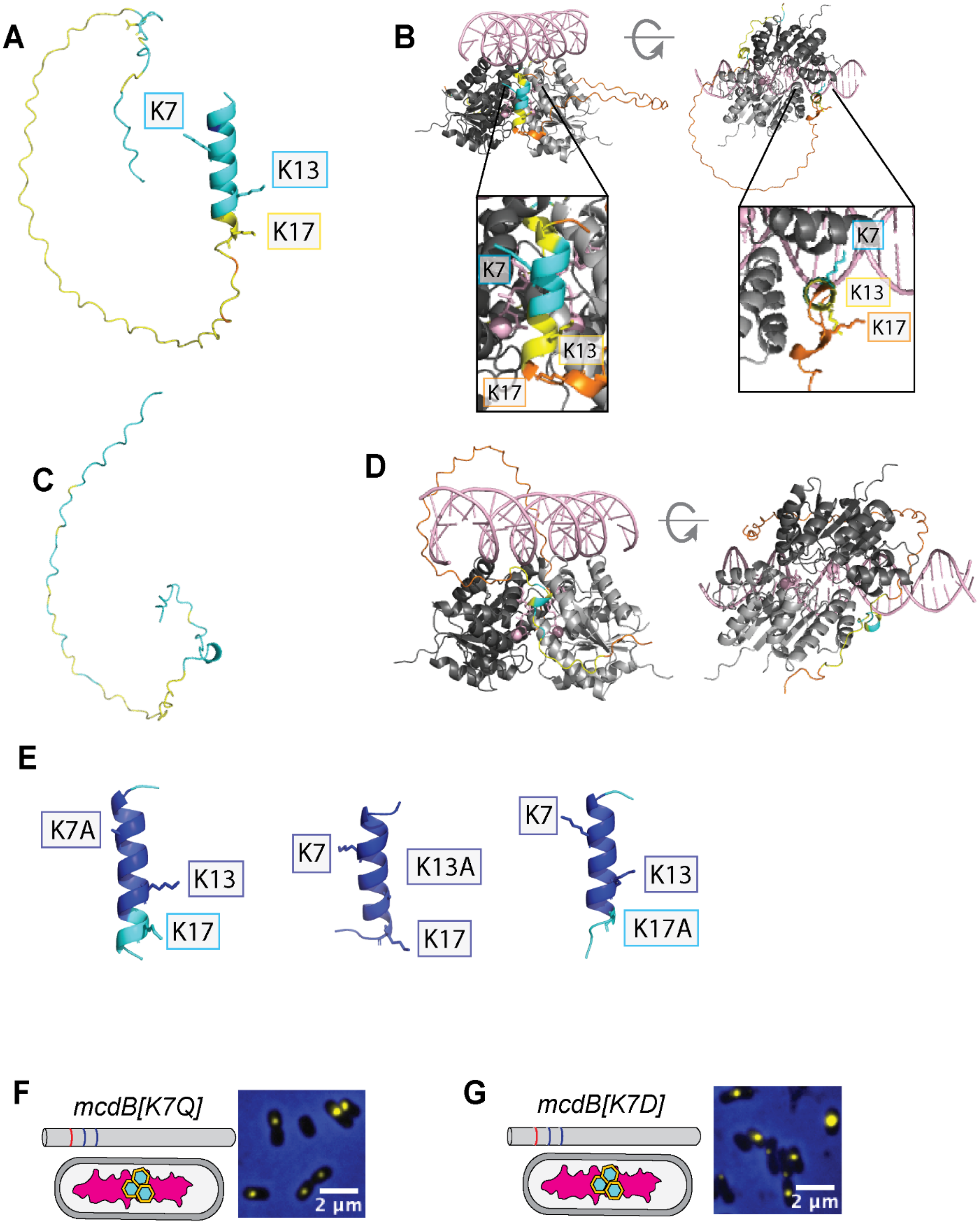

A
K7
K13
K17
B
K7
K13
K17
K7
K13
K17
C
D
E
K7A
K13
K17
K7
K13A
K17
K7
K13
K17A
F
mcdB[K7Q]
2 μm
G
mcdB[K7D]
2 μm

**Supplemental Figure 1: AlphaFold3 can reliably predict and dock structures of McdB N-terminal isoforms.** Putative McdB structures colored by pLDDT confidence score – Dark Blue = Very high confidence (>90); Light Blue = Confident (70-90); Yellow = Low (50-70); Orange = Very low (< 50). **(A)** Predicted full-length structure of McdB, which includes region of disorder. **(B)** Full-length McdB docked with ATP-bound McdA homodimer (grey) docked onto a non-specific DNA substrate (pink). ATP and magnesium ions are also pink. ipTM is 0.81 **(C)** Predicted 20 amino acid N-terminal truncation (Δ1-20) of McdB. **(D)** N-terminal truncation (Δ1-20) of McdB docked with McdA. ipTM is 0.82. **(E)** 20-amino acid N-terminal peptides of McdB [K7A], McdB [K13A], and McdB [K17A] both alone and docked with McdA. Respective ipTMs are 0.89, 0.89, and 0.88.

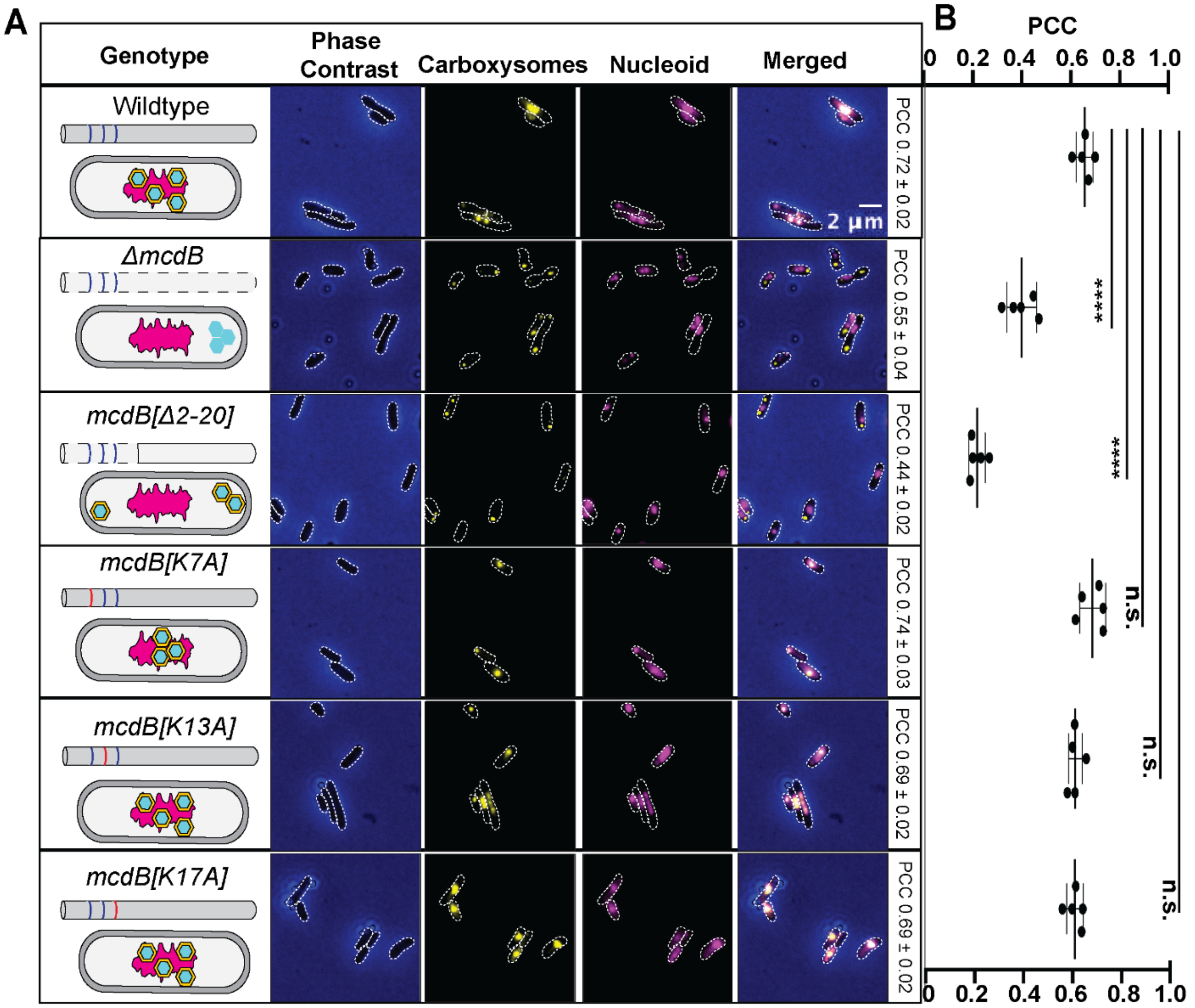


**Supplemental Figure 2: The first 20 amino acids of McdB dictate carboxysome localization relative to the nucleoid. (A)** Carboxysomes associate with DAPI-stained DNA and form mid-cell aggregates if K7 is modified to a residue that is polar and uncharged or **(B)** polar and negatively charged. **(C)** Carboxysomes remain colocalized with the DAPI-stained nucleoid in wildtype, *mcdB* [K7A], *mcdB* [K13A], and *mcdB* [K17A] cells. However, carboxysomes aggregate and become nucleoid-excluded in Δ*mcdB* and *mcdB* [Δ2-20] strains, despite the increased cytoplasmic space. Scale bar: 2 µm. **(D)** Pearson correlation coefficient (PCC) represents the positive colocalization of the bacterial chromosome and carboxysomes, 1 being the highest degree of colocalization and 0 being complete dissociation.

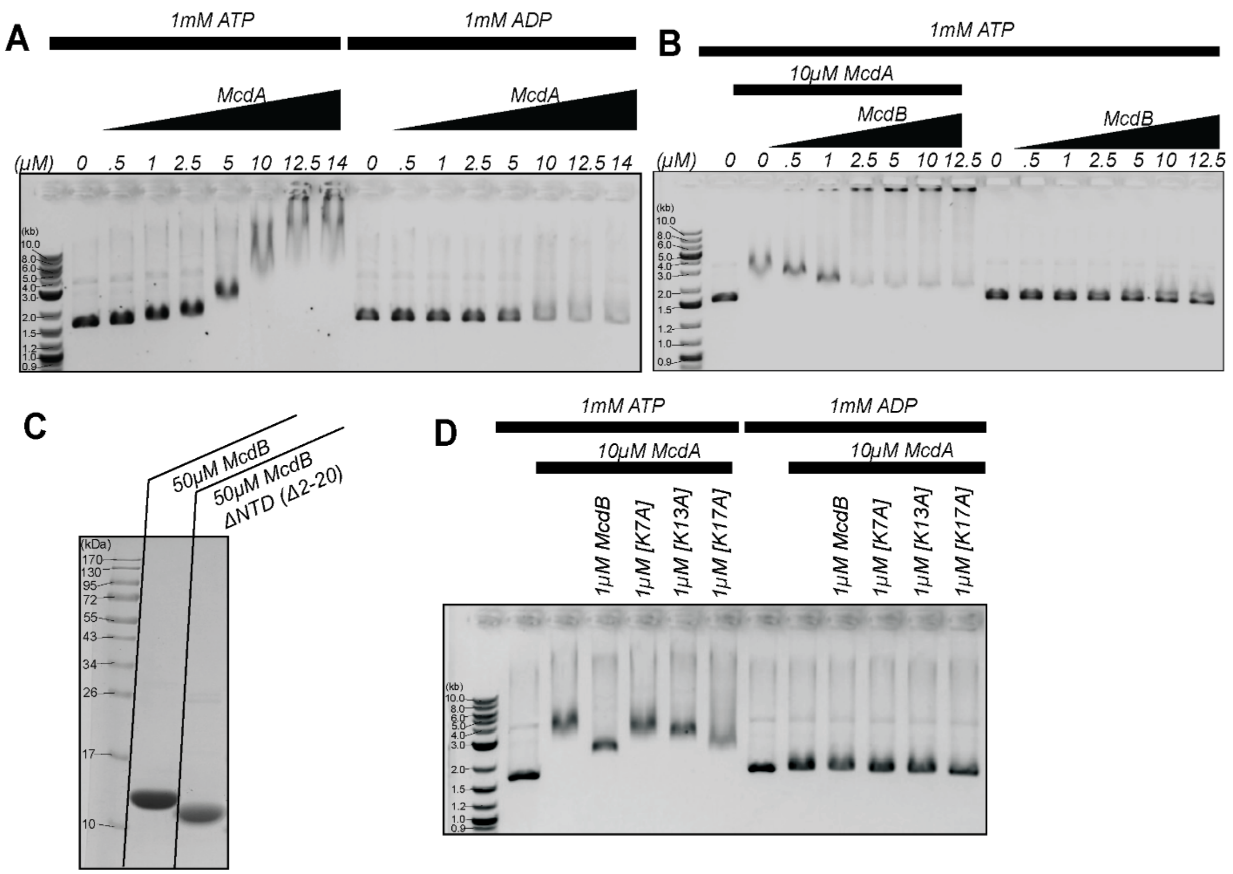


**Supplemental Figure 3: Purified McdA and McdB functionally associate in DNA-binding assays. Images are representative of three independent assays. (A)** Purified McdA forms a high-order complex (DNA-McdA) in an ATP-dependent manner. The plasmid pUC19 (2.7 kb) was incubated with increasing amounts of McdA (0, 0.5, 1, 2.5, 5, 10, 12.5, 14 μM) in the presence of either ATP or ADP. **(B)** Purified McdB forms a high-order complex in a McdA-dependent manner. Incremental addition of wildtype McdB results in large complex formation DNA:McdA:McdB. **(C)** Electrophoretic mobility shift assays show incremental release of DNA:McdA:mutant McdB complex. The addition of 1uM McdB [K7A] favors larger complex formation as compared to wildtype. McdB [K17A] most reassembles wildtype McdB, while McdB [K13A] resembles an intermediate state. Image is representative of three independent assays. **(D)** Wildtype McdB and McdB[Δ 2-20] run a single species on SDS-PAGE gel.

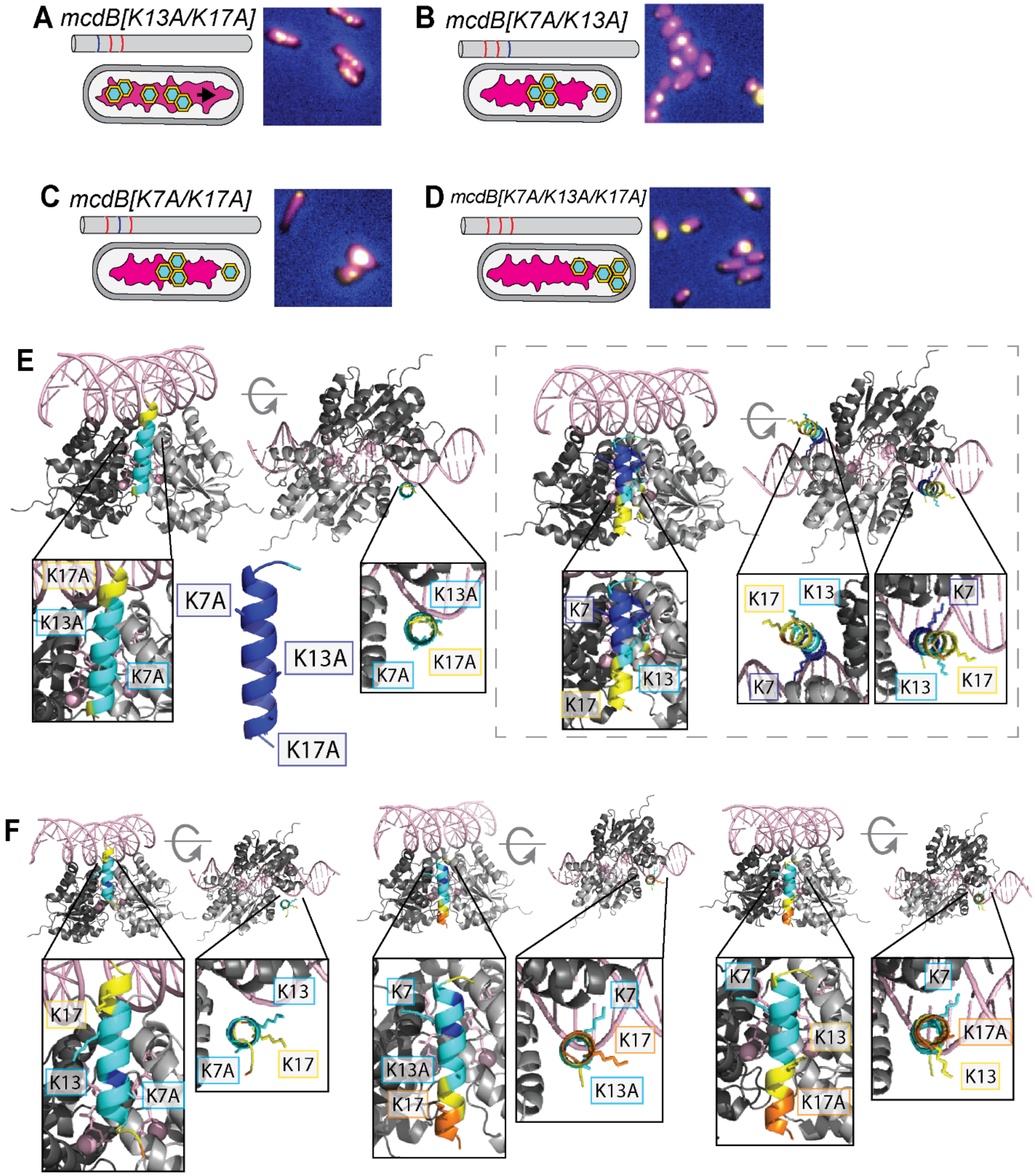
A mcdB[K13A/K17A]
B mcdB[K7A/K13A]
C mcdB[K7A/K17A]
D mcdB[K7A/K13A/K17A]
E
K17A
K13A
K7A
K7A
K13A
K17A
K13A
K7A
K17A
K7
K13
K17
K17
K13
K7
K7
K13
K17
F
K17
K13
K7A
K13
K7A
K17
K7
K13A
K17
K7
K17
K13A
K7
K13
K17A
K7
K17A
K13

**Supplemental Figure 4: McdB is still predicted to associate with McdA even when multiple N-terminal lysines are mutated. (A)** Carboxysomes are distributed across the bacterial nucleoid if K13 and K17 are both modified to a residue that is nonpolar and uncharged. **(B)** Carboxysomes form polar and mid-cell aggregates if K7 and K13 are both modified to a residue that is nonpolar and uncharged. Moderate change in cell morphology is noted. **(C)** Carboxysomes form polar and mid-cell aggregates if K7 and K17 are both modified to a residue that is nonpolar and uncharged. Change in cell morphology is noted. **(D)** Carboxysomes form polar and mid-cell aggregates if K7, K13, and K17 are all modified to a residue that is nonpolar and uncharged. Moderate heterogeneous changes in cell morphology is noted. **(E)** 20-amino acid N-terminal peptide of McdB [K7A/K13A/K17A] colored by pLDDT confidence score – Dark Blue = Very high confidence (>90); Light Blue = Confident (70-90); Yellow = Low (50-70); Orange = Very low (< 50). 2 McdA subunits form a homodimer (grey) docked with the 20-amino acid N-terminal peptide of McdB [K7A/K13A/K17A] DNA, ATP, and $Mg^{2+}$ cofactors are pink. ipTM is 0.89

**VIDEO LEGENDS:**

**Video 1 – Carboxysomes are distributed with wildtype McdB.** Carboxysomes (CbbS-mTQ, shown in yellow) are equidistantly positioned in wildtype cells. 12.5-hour video, each frame is 15 minutes (scale bar 2 μm).

**Video 2 – Carboxysomes form nucleoid-excluded aggregates at the cell pole in ΔMcdB cells.** Carboxysomes (CbbS-mTQ, shown in yellow) form polar foci. 12.5-hour video, each frame is 15 minutes (scale bar 2 μm).

**Video 3 - Carboxysomes form nucleoid-excluded aggregates at the cell pole in McdB[Δ2-20] cells.** Carboxysomes (CbbS-mTQ, shown in yellow) form polar foci. 12.5-hour video, each frame is 15 minutes (scale bar 2 μm).

**Video 4 - Carboxysomes are distributed across the nucleoid region in McdB[K13A] cells.** Carboxysomes (CbbS-mTQ, shown in yellow) are equidistantly positioned. 12.5-hour video, each frame is 15 minutes (scale bar 2 μm).

**Video 5 - Carboxysomes are distributed across the nucleoid region in McdB[K17A] cells.** Carboxysomes (CbbS-mTQ, shown in yellow) are equidistantly positioned. 12.5-hour video, each frame is 15 minutes (scale bar 2 μm).

**Video 6 - Carboxysomes cluster at mid-nucleoid in McdB[K7A] cells.** Carboxysomes (CbbS-mTQ, shown in yellow) form a focus at mid-cell. 12.5-hour video, each frame is 15 minutes (scale bar 2 μm).

**Supplemental Table 1: Strain List**

| **Species** | **Strain Number/Identifier** | **Genotype** |
|---|---|---|
| *E. coli* | 738 / pLT18 | Hn::Cbbs-mTQ △Hn0911(McdB) |
| *E. coli* | 787 / pJAB2 | Hn::CbbS-mTQ, native McdB[K7A] |
| *E. coli* | 788 / pJAB3 | Hn::CbbS-mTQ, native McdB[K13A] |
| *E. coli* | 789 / pJAB4 | Hn::CbbS-mTQ, native McdB[K17A] |
| *E. coli* | 861 / pJAB37 | Hn::CbbS-mTQ, native McdB[△2-20] |
| *H. neapolitanus* | 2 | Hn::CbbS-mTQ |
| *H. neapolitanus* | 114 | Hn::Cbbs-mTQ △Hn0911 (McdB) |
| *H. neapolitanus* | 473 | Hn::CbbS-mTQ, native McdB[K7A] |
| *H. neapolitanus* | 474 | Hn::CbbS-mTQ, native McdB[K13A] |
| *H. neapolitanus* | 475 | Hn::CbbS-mTQ, native McdB[K17A] |
| *H. neapolitanus* | 553 | Hn::CbbS-mTQ, native McdB[△2-20] |

**Supplemental Table 2: Parameter Thresholds**

| Parameter | Description | Wildtype | Mutants |
|---|---|---|---|
| *denoise_radius* | Radius for Gaussian blur in the *skimage.filters.unsharp_mask* function | 3 | 5 |
| *denoise_amount* | Scaling factor for detail amplification in the *skimage.filters.unsharp_mask* function | 20 | 20 |
| *gauss_blur* | Gaussian blur radius applied to image post denoising | 0.5 | 0.5 |
| *detect_threshold* | Intensity threshold for the *skimage.feature.blob_log* function | 0.01 | 0.05 |
| *overlap* | Parameter for the *skimage.feature.blob_log* function which defines the amount of overlap for two adjacent spots. | 0.5 | 0.1 |
| *detect_min_sigma* | Minimum radius for LoG blob detection | 1.05 | 1.05 |
| *detect_max_sigma* | Maximum radius for LoG blob detection | 1.5 | 2 |
| *dfrlmz* | Expected diameter of a diffraction limited fluorescent spot | 3 | 2 |
| *stdtol* | Tolerance on the standard deviation of the 2D Gaussian fit | 1.5 | 2 |

**Supplemental Table 3: Sample Sizes for Image Analysis**

| Sample | Rep 1 | Rep 2 | Rep 3 |
|---|---|---|---|
| WT | 889 | 593 | 1986 |
| *ΔmcdB* | 996 | 1733 | 1313 |
| *mcdB [Δ2-20]* | 1117 | 3402 | 1354 |
| *mcdB* [K7A] | 852 | 212 | 428 |
| *mcdB* [K13A] | 4060 | 2109 | 873 |
| *mcdB* [K17A] | 3258 | 1712 | 1587 |
